# Activation of SGK1.1 up-regulates the M-current in the presence of epilepsy mutations

**DOI:** 10.1101/2021.10.19.464941

**Authors:** Elva Martin-Batista, Rían W. Manville, Belinda Rivero-Pérez, David Bartolomé-Martín, Diego Alvarez de la Rosa, Geoffrey W. Abbott, Teresa Giraldez

## Abstract

In the central nervous system, the M-current plays a critical role in regulating subthreshold electrical excitability of neurons, determining their firing properties and responsiveness to synaptic input. The M-channel is mainly formed by subunits Kv7.2 and Kv7.3 that co-assemble to form a heterotetrametric channel. Mutations in Kv7.2 and Kv7.3 are associated with hyperexcitability phenotypes including benign familial neonatal epilepsy (BFNE) and neonatal epileptic encephalopathy (NEE). SGK1.1, the neuronal isoform of the serum and glucocorticoids-regulated kinase 1 (SGK1), increases M-current density in neurons, leading to reduced excitability and protection against seizures. Herein, using two-electrode voltage clamp on *Xenopus laevis* oocytes, we demonstrate that SGK1.1 selectively activates heteromeric Kv7 subunit combinations underlying the M-current. Importantly, activated SGK1.1 is able to up-regulate M-channel activity in the presence of two different epilepsy mutations found in Kv7.2 subunit, R207W and A306T. In addition, proximity ligation assays in the N2a cell line allowed us to address the effect of these mutations on Kv7-SGK1.1-Nedd4 molecular associations, a proposed pathway underlying M-channel up-regulation by SGK1.1

## INTRODUCTION

Mutations affecting ion channel subunit-encoding genes underly several different forms of epilepsy (for a recent review see (Oyrer et al. 2018)). In the CNS, *KCNQ2* and *KCNQ3* genes encode, respectively, the Kv7.2 and Kv7.3 K+ channel subunits, which co-assemble and form a heterotetrametric channel that underlies the M current (Brown and Adams 1980; Wang et al. 1998). Kv7.4 and Kv7.5 also contribute to generating the M-current in the CNS, although to a lesser extent (Kubisch et al. 1999; Lerche et al. 2000; Schroeder et al. 2000; Shah et al. 2002). M-channels are active at subthreshold membrane potentials (near −60 mV). Their activation is slow and thus they do not contribute to the repolarization of individual action potentials. M-channels do not inactivate and therefore generate a steady voltage-dependent outward current that leads to the stabilization of membrane potential, contributing to setting the resting membrane potential (RMP). The M-current is negatively regulated by activation of muscarinic acetylcholine receptors through phosphatidylinositol 4,5-bisphosphate (PIP_2_) depletion. Thus, the M-current plays a critical role in dynamically regulating subthreshold electrical excitability of neurons, determining their firing properties and responsiveness to synaptic input (Brown and Adams 1980; Constanti and Brown 1981; Wang and McKinnon 1995). Mutations in *KCNQ2* and *KCNQ3* associate with hyperexcitability phenotypes, including benign familial neonatal epilepsy (BFNE) and neonatal epileptic encephalopathy (NEE) (Singh et al. 1998; Singh et al. 2003; Soldovieri et al. 2014; Nappi et al. 2020). BNFE is a dominantly inherited condition affecting newborns and characterized by the occurrence of focal, multifocal or generalized seizures. BFNE patients have a higher risk of developing epilepsy later in life when compared to the general population (Ronen et al. 1993; Plouin and Anderson 2005). In NEE, encephalopathy is present from birth and persists during a period when seizures are uncontrolled, leading to developmental impairment. Cessation of seizures generally occurs between age nine months and four years (Miceli et al. 2010). Modulation of M-current constitutes a potential target to treat epilepsy and other diseases driven by neuronal hyperexcitability such as neuropathic pain, ischemia and schizophrenia (Wulff et al. 2009). In fact, a number of pharmacological tools have been identified which modulate M-current. Among them, retigabine and flupirtine have demonstrated ability to prevent seizure activity both in animal models and in clinical trials (Wickenden et al. 2004). However, due to drug-induced liver injury and tissue discoloration by flupirtine and retigabine, respectively, both drugs were recently discontinued (Michel et al. 2012; Clark et al. 2015; Surur et al. 2019). Since retigabine and flupirtine adverse effects are not related to their mechanism of action, it seems promising to continue studying M-current modulators and activation pathways as effective and clinically relevant anticonvulsant treatments (Surur et al. 2019). We previously described that SGK1.1, the neuronal isoform of the serum and glucocorticoid-induced kinase 1 (SGK1), up-regulates Kv7.2/3 channels by increasing their membrane abundance in a PIP_2_-dependent manner, providing a convergence point to M-channel regulation (Miranda et al. 2013). Recently, we described that M-channel up-regulation by transgenic expression of constitutively active SGK1.1 potently reduces seizure severity and duration in a kainic acid mouse model of temporal lobe epilepsy (Armas-Capote et al. 2020) and contributes to neuroprotection (Martin-Batista et al. 2021). However, the wider effects of SGK1.1 on the Kv7 family of channels forming the neuronal M-current have not been directly addressed and, most importantly, its potential role in rescuing epileptogenic mutations such as those found in Kv7.2 (R207W and A306T) is currently unknown. R207W is a mutation that neutralizes a charged amino-acid residue within the S4 transmembrane segment of the Kv7.2 voltage sensor domain, slowing voltage-sensor dependent channel activation and resulting in BFNC (Dedek et al. 2001). Mutation A306T is located in the S6 segment of Kv7.2, within the pore domain. This mutant exhibits reduced currents but retains many of the biophysical characteristics of the wild-type channel (Xiong et al. 2008). In this work, we examined the regulation by SGK1.1 of several combinations of Kv7 subunits, providing evidence that SGK1.1 regulates not only Kv7.2/3, but also Kv7.4/5, M-current forming heteromeric channels. Fully activated SGK1.1 was able to partially rescue activity in Kv7.2/3 channels harboring loss-of-function Kv7.2 epileptogenic mutations, further supporting its role as a possible avenue to counteract neuronal hyperexcitability by potentiating the M-current.

## MATERIAL AND METHODS

Animal handling and experimental procedures were approved by Universidad de La Laguna Ethics Committee and conform to Spanish and European guidelines for protection of experimental animals (RD53/2013; 2010/63/EU).

### Plasmid constructs, cRNA synthesis and oocyte microinjection

pSRC5 plasmids carrying human Kv7.2 and Kv7.3 were kindly provided by Dr. Alvaro Villarroel (CSIC Institute of Biofisika, Vizcaya, Spain). pTLN plasmids carrying human KCNQ1, Kv7.3(A315T) and KCNQ4-5 as well as KCNE1 cloned into pRAT have been previously described (Manville et al. 2018). Mouse SGK1.1 (wild type and constitutively active mutant S515D) cloned into pcDNA3.1/V5-His-TOPO (Invitrogen) were kind gifts from Dr. Cecilia Canessa (Yale University, New Haven, CT). Kinase-dead mutant K220A was described previously (Wesch et al. 2010). SGK1.1 constructs were cloned into pECFP-N1 and pEYFP-N1 (Clontech) plasmids for expression in mammalian cells (Wesch et al. 2010). pcDNA3.1 containing N-methyl-D-aspartate receptor (NMDAR) subunit 1 (NR1) fused to YFP was a gift from Dr. Stefano Vicini (Addgene plasmid #17928; (Luo et al. 2002). KCNQ2 mutants were generated by site-directed mutagenesis using a QuikChange kit according to manufacturer’s protocol (Stratagene, San Diego, CA). cRNA transcripts encoding human KCNQ1-5 were generated by *in vitro* transcription using the T7 polymerase mMessage mMachine kit (Thermo Fisher Scientific) or SP6 polymerase mMessage mMachine kit (Thermo Fisher Scientific), after vector linearization. cRNA was quantified by spectrophotometry. Oocytes were obtained from Ecocyte Bioscience (Austin, TX) or harvested from *Xenopus laevis* females and cRNAs microinjected as previously described (Miranda et al. 2013; Manville et al. 2018). Briefly, defolliculated stage V and VI oocytes were injected with Kv7 channel α subunit cRNAs (10 ng), alone or with SGK1.1 cRNA (10 ng). Oocytes were incubated at 16° C in ND96 solution (in mM: 96 NaCl, 2 KCl, 1.8 CaCl_2_, 1 MgCl_2_, 20 HEPES, pH 7.6) containing gentamycin with daily washing for 1-2 days prior to electrophysiological recordings.

### Two-electrode voltage clamp

Two electrode voltage clamp (TEVC) was performed at room temperature with an OC-725C amplifier (Warner Instruments, Hamden, CT) and pClamp8 software (Molecular Devices, Sunnyvale, CA) 1-2 days after cRNA injection as described above. Oocytes were placed in a small volume oocyte bath in recording solution (in mM: 96 NaCl, 4 KCl, 1 MgCl_2_, 1 CaCl_2_, 10 HEPES, pH 7.6) under a dissection microscope. Pipette resistance was 1-5 MΩ when filled with 3M KCl. Currents were recorded in response to pulses between −80 mV and +40 mV at 20 mV intervals from a holding potential of −70 mV, followed by a −30mV pulse to yield current-voltage relationships, current magnitude and for quantifying activation rate. TEVC data analysis was performed with Clampfit 10.6 (Molecular Devices). Half-maximal activation voltage calculated from Boltzmann equation fits to activation curves and activation constants were calculated from monoexponential fits to evoked currents, as described (Miranda et al. 2013).

### Cell culture, transfection and genome editing

Mouse neuroblastoma Neuro2A (N2a) cells were obtained from American Type Culture Collection (Manassas, VA) and maintained in DMEM supplemented with 10% FBS. Cells were transfected 24-48 hours before the experiment using Jetprime (Polyplus Transfection) following the manufacturer’s instructions. *Sgk1* knockout in N2a cells was performed using CRISPR/Cas9 induced non-homologous end joining using one plasmid system. SpCas9 and trans-activating CRISPR RNA (tracrRNA) were expressed using the pX459 plasmid (a gift from Dr. Feng Zhang, Addgene plasmid #62988,(Ran et al. 2013)), which includes a puromycin resistance cassette. Two guide RNAs (gRNA) were designed to cut exons 6 and 8 simultaneously, using the Breaking-Cas tool (Oliveros et al. 2016) and cloned into pX459 using the BbsI site. These exons were selected because they include the catalytic domain common to all isoforms of the kinase (Arteaga et al. 2008), ensuring knockout of both isoforms. Thirty-six hours after transfection, 3 μg/ml puromycin was added, and after 48 h, surviving cells were split and seeded at low density in 10 cm dishes. Single cell clones were collected using cloning cylinders and split in multi-well plates. PCR screening was performed using primers flanking exons 6-8, which produce a 681 bp amplicon from the wild type gene (F-check mSGK1: 5’ GTAAGGCTGTGTGCAGCGTA 3’ and R-check mSGK1: 5’ TCAAACCCAAACCAAGCAAT 3’). Clones that showed amplicon absence or altered size were selected to perform western blot analysis of SGK1 expression. Thus, the absence of SGK1 and SGK1.1 was confirmed, selecting an N2a clone knockout.

### Antibodies

For SGK1.1 detection we used a rabbit polyclonal antibody produced in our laboratory (Martin-Batista et al. 2021). SGK1 was detected using rabbit anti-SGK1 polyclonal antibody (Abcam, ab43606). Kv7.2 was detected with rabbit anti-Kv7.2 polyclonal antibody (Abcam, ab22897). YFP-tagged SGK1.1 and GFP-tagged Kv7 subunits were detected using mouse anti-GFP monoclonal antibody (Abcam, ab290). Nedd4-2 was detected with a rabbit polyclonal antibody from Cell Signaling (4013S). GAPDH was detected with a mouse monoclonal antibody (Abcam, 9484). Secondary antibodies conjugated to fluorophores were obtained from Thermo Fisher Scientific (Alexa 594, A-11042; Alexa-488, A-11008).

### Western Blotting

N2a cells or pools of oocytes injected with the same cRNA combinations were lysed in TENT buffer (in mM: 50 Tris-HCl, 5 EDTA, 150 NaCl, 1% Triton X-100, pH 7.4) containing protease and phosphatase inhibitors (Roche). Centrifugation supernatants were collected, mixed with Laemmli loading buffer, and heated up to 95 °C for 5 minutes. Proteins were separated in 10% polyacrylamide gels (Mini-PROTEAN TGX Stain-Free Gels, Bio-Rad) and transferred to polyvinyl difluoride membranes (Bio-Rad) for blotting with anti-Kv7.2 and anti-GADPH antibodies referenced above. Chemiluminescence signals were quantified using Image Lab® software 6.0 (Bio-Rad).

### Proximity Ligation Assay

Cells grown on coverslips were transiently transfected with plasmids encoding the combinations of indicated variants of SGK1.1, Nedd4-2, Kv7.2 and Kv7.3, 24 hours prior to the assay, using a JetPrime® transfection kit. After fixation with 4% formaldehyde, cells were processed for the proximity ligation assay (PLA) to detect physical proximity of different protein pairs expressed in *Sgk1*-KO N2a cells following the manufacturer’s protocol (Duolink, Olink Biosciences) and using the antibodies indicated in each case. Cells were then mounted, and images obtained using a confocal microscope (Lecia SP8), 40x magnification and a zoom factor of 2. Images were analyzed using the software provided by the manufacturer (Duolink Image Tool). Results are expressed as average number of puncta/cell area normalized to control conditions obtained using non-interacting proteins (cells expressing NR1 and SGK1.1) and compared to our reference condition of cells only transfected with the channel subunits, Kv7.2 and Kv7.3.

## RESULTS

### SGK1.1 up-regulates heteromeric Kv7 chanels underlying the neuronal M-current

Expression of wild type (WT) SGK1.1 resulted in significantly increased Kv7.2/3-mediated peak and tail currents (Fig.1A-C) without altering normalized conductance (Fig. 1D), as we have previously described (Miranda et al. 2013). Co-expression of constitutively active SGK1.1 mutant S515D activated Kv7.2/3-mediated currents to the same extent as WT SGK1.1 (Fig. 1A-C), indicating that endogenous activation of the kinase is enough to maximize its effects on the channel. Similarly, oocytes expressing heteromeric Kv7.3/5, which is a described combination associated to M-current (Schroeder et al. 2000), showed currents that were increased by SGK1.1. Tail current was significantly increased in presence of the kinase at 0, +20 and +40 mV (Fig. 2A, C). Heterologous expression of Kv7.4 or Kv7.5 resulted in K^+^ currents smaller than those elicited by heteromeric Kv7.2/3/5 (Fig. S1). In contrast to the effect observed with heteromeric channels, co-expression of SGK1.1 produced an inhibitory effect on the tail current elicited by Kv7.4 (Fig. S1C) whereas no alteration was observed in Kv7.5-mediated currents (Fig. S1E-H). When we co-expressed Kv7.1 and accessory subunit KCNE1, characteristic slow activating, voltage-dependent current reminiscent of the slow cardiac K^+^ repolarizing current (I_Ks_) was elicted (Fig. S2A). Co-expression of SGK1.1 affected neither amplitude nor conductance (Fig.S 2B-D) of the Kv7.1-KCNE1 currents.

**Figure 1.**
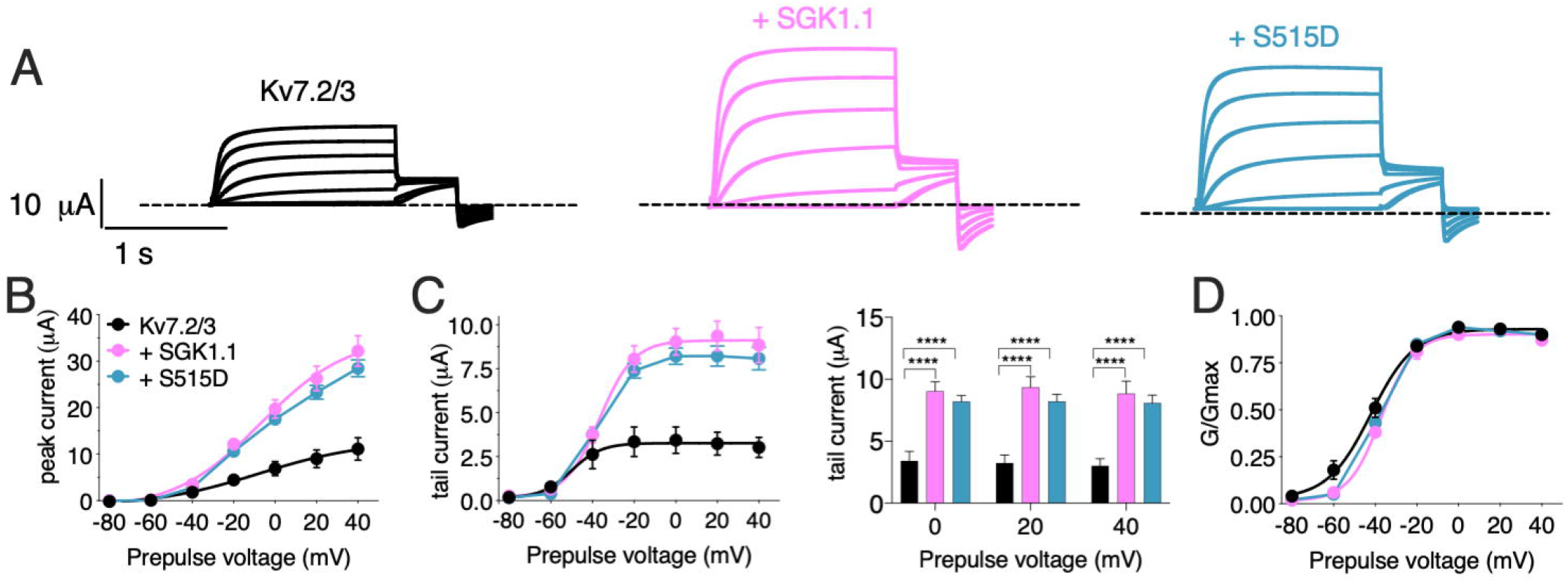
Wild-type and activated SGK1.1 up-regulate M-current. (A) Currents elicited in *Xenopus* oocytes after coinjection of cDNAs from Kv7.2/3 channel alone (first panel) or in combination with wild-type (second panel) or constitutively active (third panel) SGK1.1. (B) Peak current/voltage relationship. (C) Tail current (left) and tail currents measured at −30 mV after 0, +20 or +40 mV depolarizing pulses for the indicated construct combinations. Values represent mean ± SEM. Two-way ANOVA followed by Tukey’s correction for multiple comparisons, ****p<0.0001. (D) Normalized conductance. Legends are indicated on panel B.

**Figure 2.**
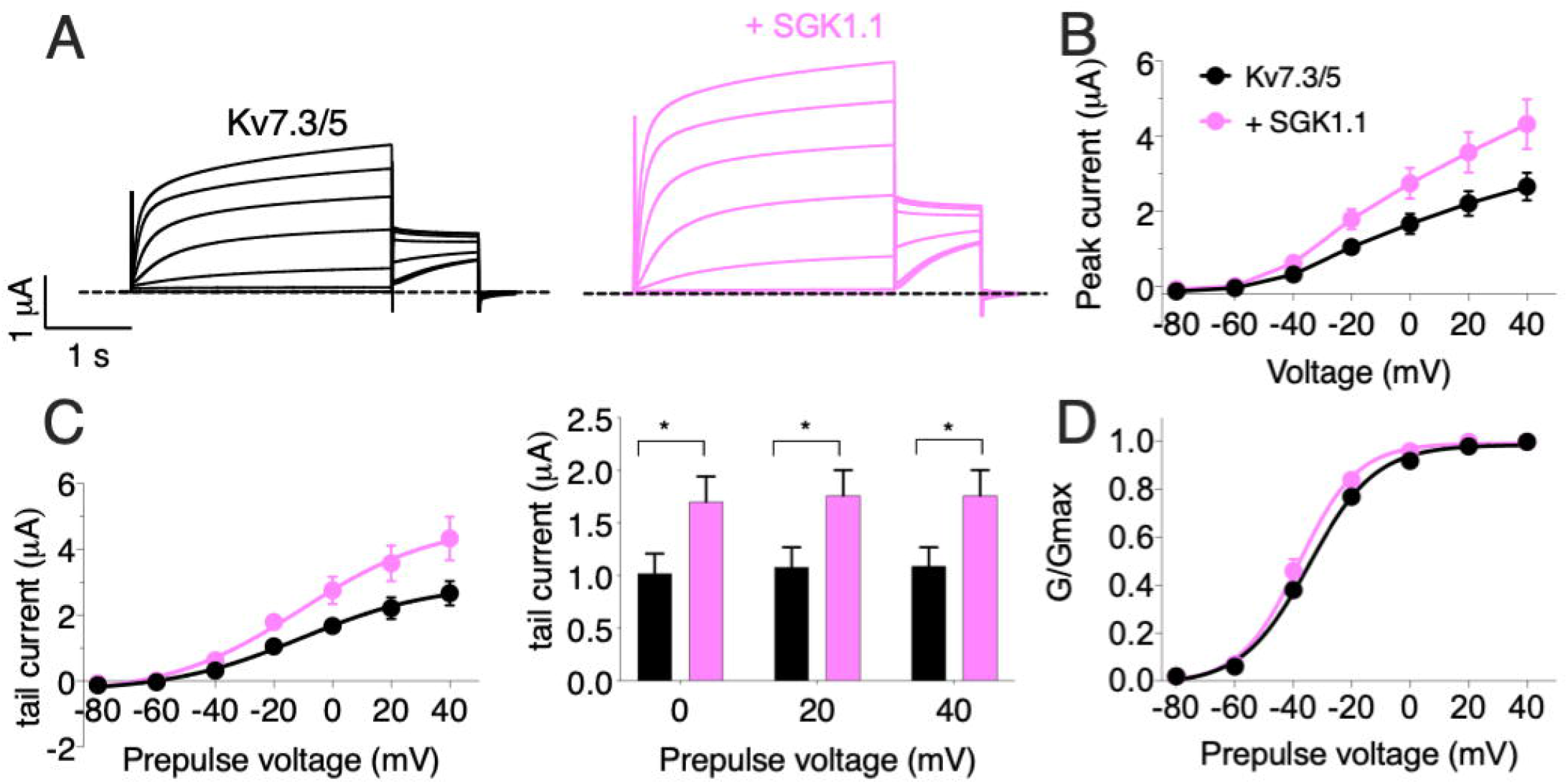
SGK1.1 increases heteromeric Kv7.3/5 currents like heteromeric Kv7.2/3, supporting the need for heteromeric channel assembly to observe SGK1.1 effects. (A) Currents elicited in Xenopus oocytes after coinjection of cRNAs from Kv7.3/5 channel alone (first panel) or in combination with wild-type SGK1.1 (second panel). (B) Peak current/voltage relationship. (C) Tail current (left) and tail currents measured at −30 mV after 0, +20 or +40 mV depolarizing pulses for the indicated construct combinations (right). Values represent mean ± SEM (Multiple t-test Holm-Sidak correction method;*p<0.05). (D) Normalized conducance. Legends are indicated on panel B.

### Activated SGK1.1 up-regulates Kv7.2 epilepsy mutant R207W and A306T channel activity in heteromeric assembly with Kv7.3

Expression of epilepsy mutant Kv7.2(R207W) (Fig. 3A) along with Kv7.3 produced smaller currents with slower voltage-dependent activation profile (Fig. 3B-D, activation constants 1006 ms (R207W) vs 104 ms (WT)), in agreement with previous reports (Dedek et al. 2001). Co-expression of wild type SGK1.1 did not produce any change in the amplitude of the R207W-modified M-current. Strikingly, however, SGK1.1 (S515D) up-regulated Kv7.2(R207W)/WT Kv7.3 peak and tail currents significantly (Fig. 3B-D). Mutation A306T, located within the pore domain in the S6 segment of the subunit (Fig. 3A), maintains most Kv7.2 biophysical characteristics but produces significantly reduced currents (Xiong et al. 2008). Co-expression of Kv7.2(A306T) with Kv7.3 did not induce detectable currents (data not shown). Therefore, we used a previously described mutation (A315T) in the pore of Kv7.3 that greatly increases current amplitude (Gomez-Posada et al. 2011). We have previously demonstrated that SGK1.1 does not affect currents elicited by Kv7.3(A315T), but does increase those of the heteromeric Kv7.2/3(A315T) channel (Miranda et al. 2013). When analyzing the currents elicited by mutant Kv7.2(A306T) in heteromeric assembly with Kv7.3(A315T) we observed similar findings to those with Kv7.2(R207W). Only the expression of the SGK1.1(S515D) mutant resulted in substantial increase of the current (Fig. 4A-C), with negligible effects on voltage dependence of activation (Fig. 4D).

**Figure 3.**
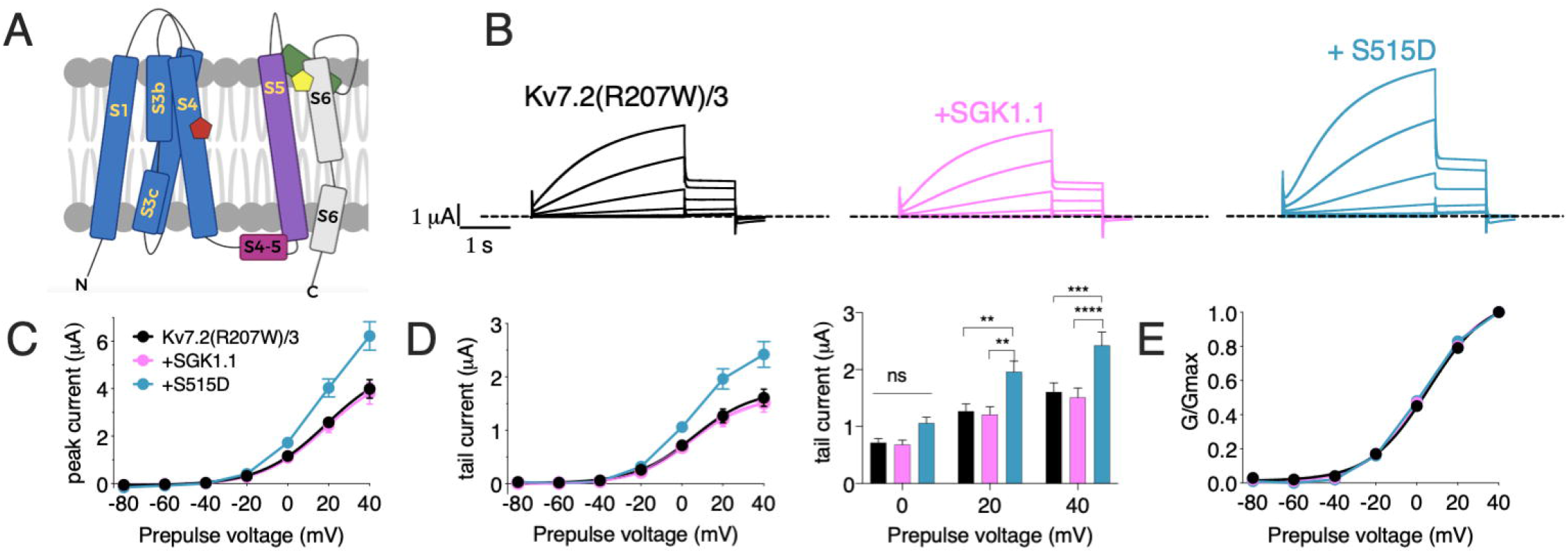
Constitutively active form SGK1.1(S515D) up-regulates Kv7.2 epilepsy mutation R207W in heteromeric assembly with Kv7.3 while WT SGK1.1 fails to. (A) Schematic representation of Kv7 channel structure. The basic organization of potassium channels is a tetramer with each monomer containing one pore-forming domain (PD) (transmembrane segments S1-S4) and a voltage sensor domain (VSD) (transmembrane segments S5-S6). Epilepsy mutants are represented as colored pentagons (R207W in red and A306T in yellow). (B) Currents elicited in Xenopus oocytes after coinjection of cDNAs from Kv7.2(R207W)/3 channel alone (first panel) or in combination with wild-type (second panel) or constitutively active (third panel) SGK1.1. (C) Peak current/voltage relationship. (D) Tail current (left) and tail currents measured at −30 mV after 0, +20 or +40 mV depolarizing pulses for the indicated construct combinations (right). Values represent mean ± SEM (Two-way ANOVA, Tukey’s correction for multiple comparisons; ns, not significant; **p<0.01; ***p<0.0005; ****p<0.0001). (E) Normalized conductance. Legends are indicated on panel C.

**Figure 4.**
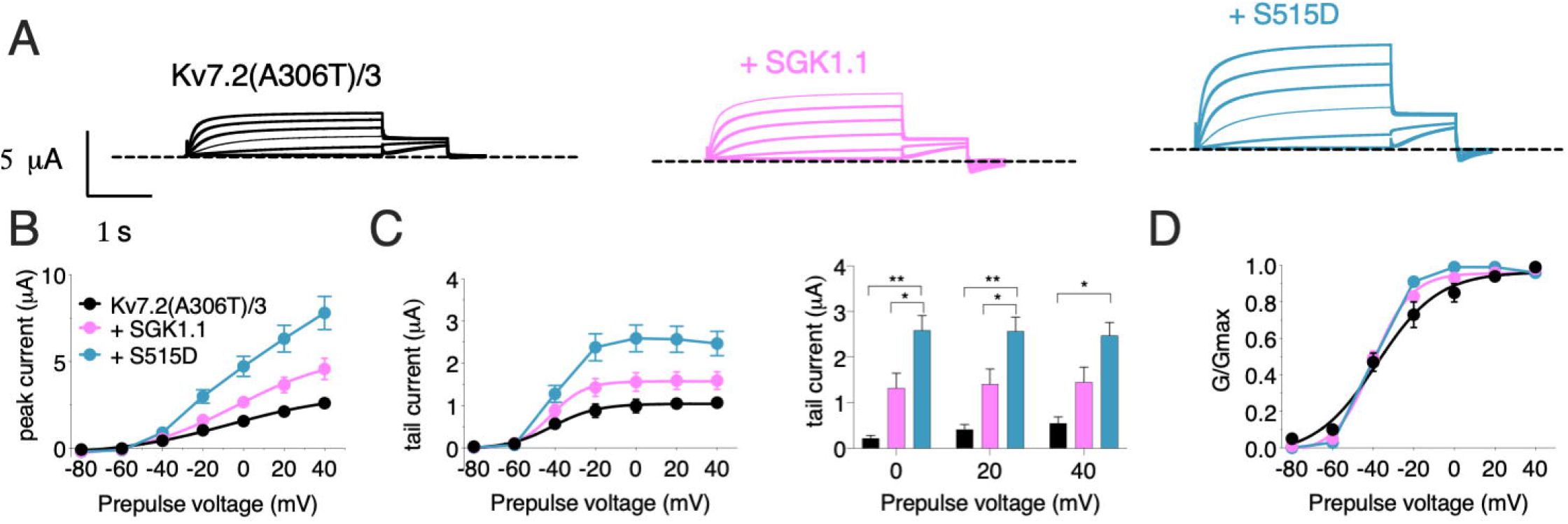
Constitutively active form of SGK1.1(S515D) up-regulates Kv7.2 epilepsy mutation A306T in heteromeric assembly with Kv7.3(A315T). (A) Currents elicited in Xenopus oocytes after coinjection of cDNAs from Kv7.2(A306T)/3 channel alone (first panel) or in combination with wild-type (second panel) or constitutively active (third panel) SGK1.1. (B) Peak current/voltage relationship. (C) Tail current (left) and tail currents measured at −30 mV after 0, +20 or +40 mV depolarizing pulses for the indicated construct combinations (right). Values represent mean ± SEM (Two-way ANOVA Tukey’s correction for multiple comparisons; *p<0.05; **p<0.01). (D) Normalized conductance. Legends are indicated on graph B.

### SGK1.1 is associated to Kv7.2/3 channels and Nedd4-2

To gain further information about the effects of these mutations, we next tested their effects on the levels of protein expression of Kv7.2 in both heterologous expression systems - injected oocytes and transfected N2a cells. Mutations R207W and A306T each significantly decreased the abundance of Kv7.2 protein in oocytes. Neither WT SGK1.1 nor activated SGK1.1(S515D) increased its expression (Fig. 5A-B). Similar results were observed in N2a (Fig. 5C-D). We previously demonstrated that SGK1.1 up-regulates Kv7.2/3 current by increasing channel membrane abundance through a Nedd4-2-mediated pathway (Miranda et al. 2013). To further determine whether these proteins form stable complexes and how epilepsy mutations in Kv7.2 may disrupt them, we performed experiments using the proximity ligation assay (PLA) in mouse neuroblastoma N2a cells with genetic inactivation of all endogenous *Sgk1* isoforms (Fig. 5E). Expression of the transfected constructs and the absence of endogenous SGK1.1 were confirmed by immunocytochemistry (Fig. S3). PLA signals were significantly more numerous in cells expressing Kv7.2 or Kv7.3 along with activated SGK1.1(S515D) compared to the reference control only expressing Kv7.2/3 (Fig. 5-F,G). Coexpression of heteromeric Kv7.2/3 significantly increased SGK1.1 association to the channel compared to homomeric Kv7.3. Differences in signal strength between homomeric and heteromeric structures suggest that association of SGK1.1(S515D) with the channel becomes facilitated in the heteromeric configuration, which supports previous electrophysiological data showing that the SGK1.1 effect requires the heteromeric assembly of Kv7.2/3 channels (Miranda et al. 2013). Association between the kinase and the heteromeric channel was significantly impaired in the presence of K220A, a mutation in the ATP-binding cassette of the protein that abolishes the kinase activity (Wesch et al. 2010). Molecular proximity between the kinase and the heteromeric channels containing epileptogenic mutations was detectable (Fig. 5F,G), even if the mutant subunits are expressed at lower levels.

**Figure 5.**
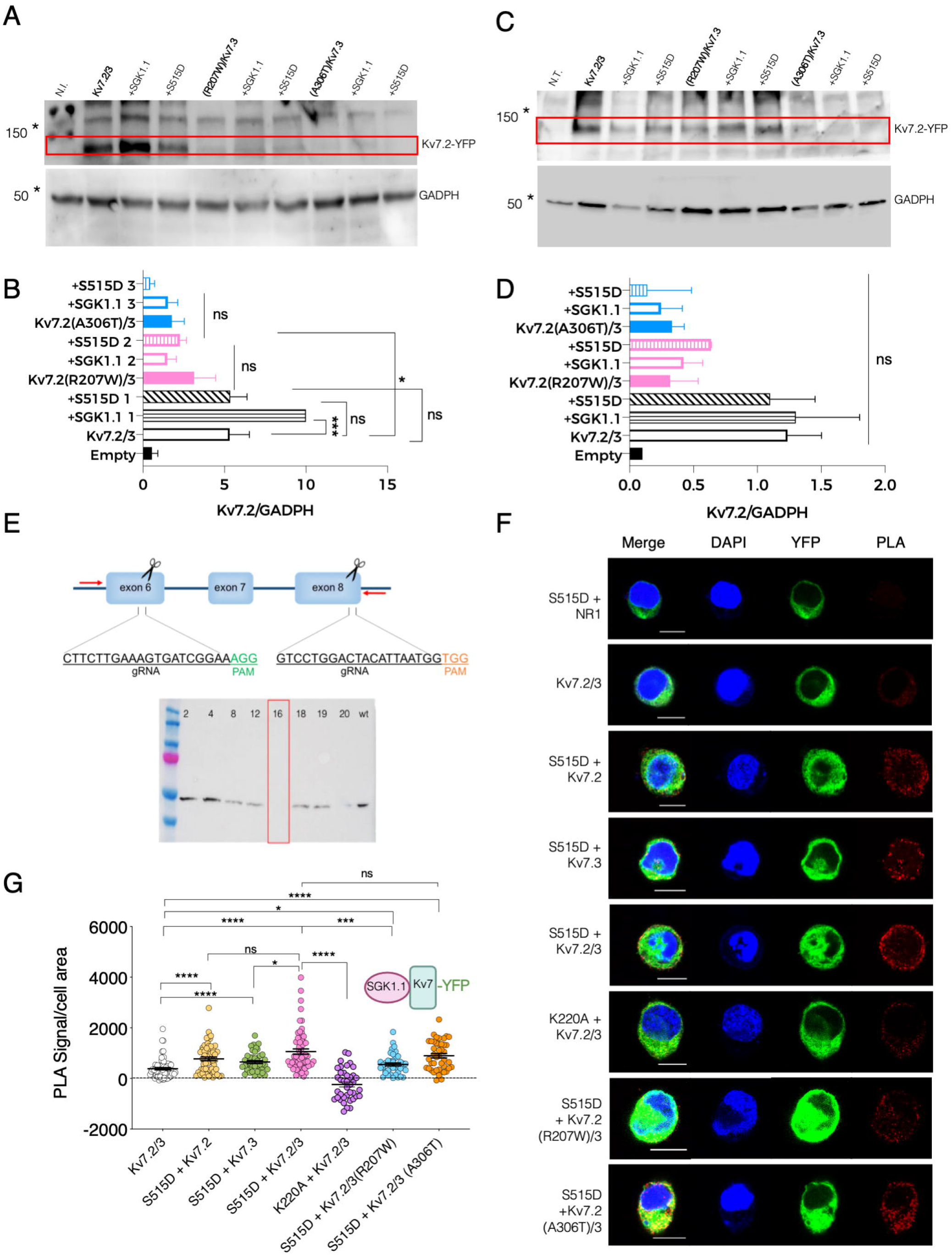
Association between SGK1.1(S515D) and heteromeric channel Kv7.2/3 is reduced in presence of epilepsy mutant Kv7.2(R207W) but not Kv7.2(A306T). (A) Representative immunoblot showing expression levels of Kv7.2 fused to YFP (top panel, 110 KDa) and GADPH (bottom panel, 50 KDa) from oocytes injected with the indicated constructs; N.I., water-injected oocytes. Asterisks denote migration of the indicated molecular mass marker. (B) Quantitative analysis of Kv7.2 expression levels in *Xenopus* oocytes. Values are mean ± SEM from at least three independent experiments (ANOVA Sidak’s test for multiple comparisons; ns, not significant; *p<0.05; ***p<0.0005). (C) Representative western blot showing expression levels of Kv7.2 fused to YFP (top panel, 110 KDa approximately) and GADPH (bottom panel, 50 KDa) from N2a *Sgk1*-KO cells transfected with the indicated constructs; N.T., non-transfected cells. Asterisks denote migration of the indicated molecular mass marker. (D) Quantitative analysis of Kv7.2 expression levels in N2a cells from two independent replicates (ANOVA Sidak’s test for multiple comparisons; ns, not significant). (E) Top, schematic representation of the CRISPR/Cas9 strategy to knockout the *Sgk1* gene. Insets indicate the sequence and hybridization sites for the guides, and PAM sequences (NGG) required for Cas9 to cut the DNA at the end of each targeted exon. Screening PCR primers are indicated as red arrows. Bottom, western blot detecting the presence of SGK1 protein expression in WT N2a cells or in selected single cell clones. Clone 16 was selected and used as *Sgk1* knockout in this study. (F) PLA was performed on N2a *Sgk1*-KO cells transfected with the indicated constructs. Cells transfected with SGK1.1(S515D) and NR1 were used as negative control (see panel F and dotted line at 0 in panel G). Bars correspond to 10 μM. (G) Quantification of PLA positive signals for each condition (ANOVA Kruskal Wallis test, ns, not significant; *p<0.05; ***p<0.0005 ****p<0.0001). Each dot represents an individual cell from at least three independent experiments.

The ubiquitous isoform SGK1 inactivates Nedd4-2 by phosphorylating it at residue S448 (Debonneville et al. 2001). We previously demonstrated that SGK1.1 also phosphorylates Nedd4-2 (Armas-Capote et al. 2020), and that SGK1.1 up-regulates Kv7.2/3 currents via a Nedd4-2-mediated mechanism (Miranda et al. 2013). In this work, we used the PLA approach to determine the existence of stable associations between Nedd4-2 and SGK1.1 (S515D). Our results showed that activated SGK1.1 associates with Nedd4-2, in agreement with previous reports suggesting a direct interaction between Kv7 and Nedd4-2 (Ekberg et al. 2007), and that this association is significantly reduced in the presence of the kinase dead mutant SGK1.1 (K220A) (Fig. 6A,B).

**Figure 6.**
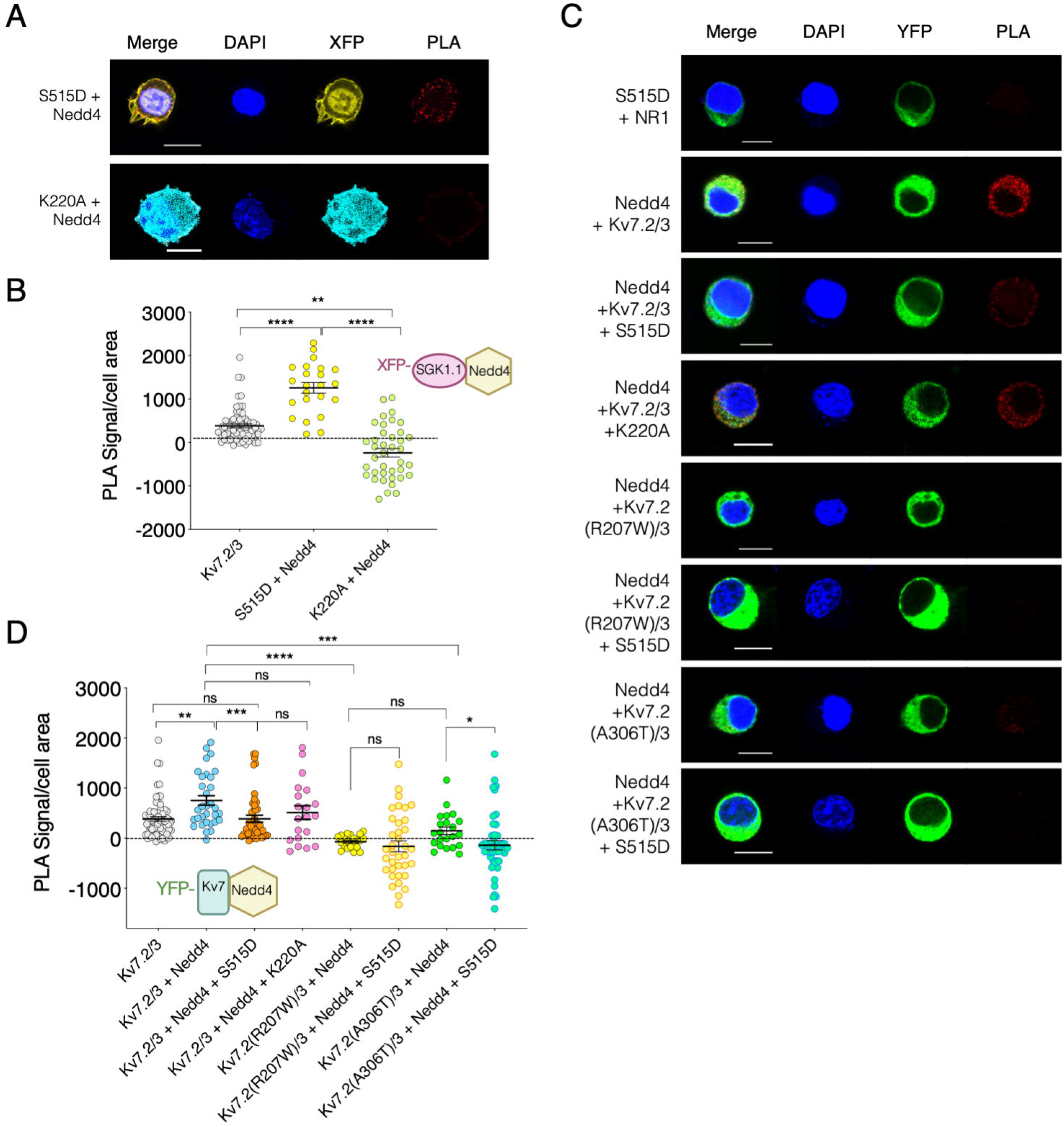
Kv7.2/3 heteromeric channel is located in close proximity to Nedd4-2 and this association weakens in presence of activated SGK1.1. (A-C) Representative images of PLA analysis performed on N2a *Sgk1*-KO cells transfected with the indicated constructs. Cells transfected with SGK1.1 (S515D) and NR1 were used as negative control (dotted line at 0 in panels B and D). Bars correspond to 10 μM. (B-D) Quantification of PLA positive signals for each condition (ANOVA Kruskal Wallis test, ns, not significant; *p<0.05; **p<0.01; ***p<0.0005 ****p<0.0001). Each dot represents an individual cell from at least three independent experiments.

We next studied the association between Nedd4-2 and Kv7 channels and whether SGK1.1 co-expression alters this complex. Our results show significantly augmented PLA signals in cells co-expressing Kv7.2/3 and Nedd4-2 as compared with the control (Fig. 6C), consistent with the association of both proteins. Importantly, this interaction was significantly reduced when the activated form of SGK1.1 was co-expressed, but not in the presence of SGK1.1 (K220A) (Fig. 6D) demonstrating that the activity of the kinase disrupts the association between Nedd4-2 and the channel. Importantly, association between the channel and the ubiquitin ligase was significantly diminished in the presence of either of the epileptogenic Kv7.2 mutations.

## DISCUSSION

The results presented here, in agreement with our previously published study (Miranda et al. 2013), demonstrate that the neuronal isoform SGK1.1 selectively upregulates heteromeric Kv7.2/3 and Kv7.3/5 channel activity, while it fails to regulate or even decreases the activity of homomeric channels formed by Kv7.2 or 7.3 (Miranda et al. 2013), Kv7.4 or 7.5. In addition, SGK1.1 was not able to modulate the current elicited by Kv7.1/KCNE1, a subunit combination that generates the major repolarizing cardiac current, I_Ks_. In contrast, the ubiquitous isoform SGK1 has been described to increase I_Ks_ currents in *Xenopus laevis* oocytes in a Nedd4-2 dependent manner (Seebohm et al. 2008). This leads us to hypothesize that the differential NH_2_-terminal domain present in SGK1.1 and its ability to interact with PIP_2_ could be responsible for the selectivity of the neuronal isoform of the kinase towards heteromeric channels underlying the M-current. Similar to Kv7.2 and Kv7.3, Kv7.5 is expressed in the brain and yields currents that form heteromeric channels with Kv7.3 and are inhibited by M1 muscarinic receptor activation (Schroeder et al. 2000). Our results demonstrate the ability of SGK1.1 to upregulate Kv7.3/5-elicited M-currents, an effect that could contribute to the anticonvulsant activity of the kinase (Miranda et al. 2013; Armas-Capote et al. 2020). In fact, some studies demonstrating activation of Kv7.3/5 by retigabine suggested it might constitute a molecular target for this agent along with Kv7.2/3 (Wickenden et al. 2001). In addition, SGK1.1 is mainly expressed in pyramidal neurons (Wesch et al. 2010; Martin-Batista et al. 2021). Altogether, these data indicate that SGK1.1 activation could be a promising therapeutic target for epilepsy, as it shares the mechanism of action of retigabine without substantial risk of activating potassium channels expressed outside neuronal tissue.

As previously mentioned, among the different causes for epilepsy, genetic mutations affecting the functionality of Kv7.2-5 channels have been described in diverse forms of the disease. We have assessed the power of SGK1.1 to upregulate the M-current in the presence of two epilepsy-associated Kv7.2 mutations. Our results suggest that constitutive activation of the kinase, which does not have an additional effect on WT Kv7.2/3-generated M-current, is able to significantly upregulate epileptogenic mutations Kv7.2(R207W) and Kv7.2(A306T). Mutation R207W, which results in BFNC and myokymia (Dedek et al. 2001), neutralizes a charged amino acid residue within the S4 segment, affecting the voltage sensor domain and slowing voltage-sensor dependent activation. Kv7.2(A306T) is located in the S6 segment (pore domain) and also associated with BFNC. In this study, we show that both mutations also result in a reduction in protein expression levels, at least in heterologous expression systems. Thus, our findings might contribute to understand the phenotype observed in patients carrying these mutations because, other than being affected by the intrinsic malfunction of the mutated channel, we can now suggest that the expression of the protein and the ability of wild type SGK1.1 to up-regulate it are disturbed. Therefore, activated SGK1.1 could constitute a strategy to increase the M-current in the presence of mutations that diminish this potassium current. Whether or not pharmacological activation of SGK1.1 may provide a useful therapeutic approach should be further addressed.

As part of our interest in understanding the effect of epileptogenic Kv7.2 mutations on the SGK1.1 mechanism of action, we evaluated the physical proximity between the kinase and the channel by PLA. Our results demonstrate that the association between SGK1.1 and heteromeric Kv7.2/3 was significantly higher than that observed in the presence of homomeric Kv7.3 channels, in agreement with the functional data, which indicate that the SGK1.1 effect requires the heteromeric co-assembly of Kv7.2/3 (Miranda et al. 2013) or Kv7.3/5 channels. The fact that SGK1.1 still associates with, but fails to regulate, homomeric channels supports the hypothesis that the mechanism controlling their plasma membrane levels is different from the one regulating heteromeric combinations of Kv7 subunits. Association of SGK1.1 with the heteromeric channel was significantly impaired in the inactive mutant SGK1.1 (K220A), suggesting that conformational changes associated to the catalytic activity of the kinase are crucial for its interaction with the channel. In addition, we were able to detect association of Nedd4-2 with SGK1.1 and, most importantly, to Kv7 channels in a SGK1.1-dependent manner. As we have previously demonstrated, SGK1.1 enhances the levels of phosphorylation of Nedd4-2 at residue S448 (Armas-Capote et al. 2020), similarly to SGK1, a process that results in the repression of Nedd4-2 ubiquitylation activity (Debonneville et al. 2001). This may prevent M-channel degradation, stabilizing it in the membrane. Consistently, we found that constitutively active, but not the kinase-dead mutant of SGK1.1 is able to displace Nedd4-2 from its interaction with the M-channel.

Co-expression of SGK1.1 with Kv7.2 epilepsy mutants produced divergent results. Both mutants were still able to closely associated with SGK1.1, even though they are expressed at significantly reduced levels. Therefore, it appears that overall Kv7 subunit abundance is not a predictor of SGK1.1 association, which may be compartment-specific and depend on the relative abundance of the channel in the plasma membrane vs. intracellular compartments. In contrast, association between Nedd4-2 and Kv7 was significantly reduced in the presence of either epilepsy mutation. Given that both mutations affect different functional regions of the channel, it is unlikely that they directly control the ability of Kv7.2 to interact with Nedd4-2. We speculate that their presence indirectly alters the ability of Nedd4-2 to interact with Kv7.2, possibly by altering its subcellular localization and/or expression levels. In summary, it seems clear that SGK1.1 can still interact with the M-channel independently of the presence of Nedd4-2 in the complex and regardless of mutations altering Kv7.2 activity/expression. Thus, it is tempting to speculate that activation of the kinase might be up-regulating the M-current through an alternative mechanism in addition to the Nedd4-2 pathway. This mechanism would be more prominent in the case of the epileptogenic mutations, since these mutants lack the interaction with Nedd4-2 and therefore are not subject to the same regulatory pathway as the WT channel. One possibility for such alternative pathway of regulation is phosphorylation. As has been proposed for SGK1 (Bongiorno et al. 2011), SGK1.1 could interact directly with the RXRXXS/T consensus motif in the subunits of ion channels, additionally to the indirect association through the PY motif present in Nedd4-2. The Kv7 subunit sequence shows interaction sites for phosphorylation on serine, threonine and tyrosine residues (Ismailov and Benos 1995). Even though the consensus phosphorylation motif of SGK1 has only been described in Kv7.4 (Seebohm et al. 2005), the cytoplasmic N-terminal domain in Kv7.2 contains a consensus site for cAMP-dependent phosphorylation by PKA that is required for its stimulation by cAMP (Schroeder et al. 1998). PKA, whose consensus sequence overlaps with that of SGK1, not only phosphorylates Kv7 but also Nedd4-2 at the same sites as does SGK1. Hence it would be possible that SGK1.1 phosphorylates the channel likewise (Snyder et al. 2004). In addition, mass spectrometry studies have revealed different phosphorylation sites in the S4-S5 loop of Kv7.2/3 within a sequence highly conserved among Kv7 family members (Surti et al. 2005). Whether SGK1.1 is able to directly phosphorylate Kv7 channel subunits remains unexplored. Alternatively, the kinase might be indirectly regulating channel trafficking. For instance, it has been previously demonstrated that the ubiquitous isoform SGK1 regulates Kv7.1/KCNE1 heteromer trafficking via Rab11-mediated recycling (Seebohm et al. 2008). It is clear from our data that the interaction between SGK1.1, Nedd4-2 and Kv7 α subunits is complex and depends not only on total protein abundance, but also on enzymatic activity, subcellular localization and the presence or absence of certain epilepsy mutations.

In summary, our study demonstrates that the neuronal isoform SGK1.1 selectively up-regulates Kv7 subunit heteromers underlying the M-current and that activation of this kinase may provide a therapeutic target for treating epilepsy, particularly in patients carrying specific Kv7.2 epileptogenic mutations.

## Author contributions

**EMB:** conceptualization, investigation, formal analysis, writing-original draft, writing-review & editing, visualization. **RWM:** conceptualization, investigation, formal analysis, writing-review. **BRP:** conceptualization, investigation, formal analysis, writing-review. **DBM:** conceptualization, investigation, formal analysis, writing-review. **DAdlR:** conceptualization, formal analysis, writing-original draft, writing-review & editing, visualization. **GWA:** conceptualization, formal analysis, writing-review & editing, funding acquisition, supervision, project administration. **TG:** conceptualization, formal analysis, writing-original draft, writing-review & editing, funding acquisition, supervision, visualization, project administration, validation.

## REFERENCES

Armas-Capote N, Maglio LE, Pérez-Atencio L, Martin-Batista E, Reboreda A, Barios JA, Hernandez G, Alvarez de la Rosa D, Lamas JA, Barrio LC, et al. 2020. SGK1.1 Reduces Kainic Acid-Induced Seizure Severity and Leads to Rapid Termination of Seizures. Cereb. Cortex 30:3184–3197. doi:10.1093/cercor/bhz302.

Bongiorno D, Schuetz F, Poronnik P, Adams DJ. 2011. Regulation of voltage-gated ion channels in excitable cells by the ubiquitin ligases Nedd4 and Nedd4-2. Channels 5:79–88. doi:10.4161/chan.5.1.13967.

Brown DA, Adams P. 1980. Summary for Policymakers. In: Intergovernmental Panel on Climate Change, editor. Climate Change 2013 - The Physical Science Basis. Vol. 283. Cambridge: Cambridge University Press. p. 1–30.

Clark S, Antell A, Kaufman K. 2015. New antiepileptic medication linked to blue discoloration of the skin and eyes. Ther. Adv. Drug Saf. 6:15–19. doi: 10.1177/2042098614560736.

Constanti A, Brown DA. 1981. M - current sin voltage - clamped mammalian sympathetic neurones. 24:289–294.

Debonneville C, Flores SY, Kamynina E, Plant PJ, Tauxe C, Thomas MA, Munster C, Chraibi A, Pratt JH, Horisberger JD, et al. 2001. Phosphorylation of Nedd4-2 by Sgk1 regulates epithelial Na(+) channel cell surface expression. EMBO J. 20:7052–7059. doi:10.1093/emboj/20.24.7052.

Dedek K, Kunath B, Kananura C, Reuner U, Jentsch TJ, Steinlein OK. 2001. Myokymia and neonatal epilepsy caused by a mutation in the voltage sensor of the KCNQ2 K+ channel. Proc. Natl. Acad. Sci. 98:12272–12277. doi:10.1073/pnas.211431298.

Ekberg J, Schuetz F, Boase NA, Conroy SJ, Manning J, Kumar S, Poronnik P, Adams DJ. 2007. Regulation of the voltage-gated K+ channels KCNQ2/3 and KCNQ3/5 by ubiquitination: Novel role for Nedd4-2. J. Biol. Chem. 282:12135–12142. doi:10.1074/jbc.M609385200.

Gomez-Posada JC, Aivar P, Alberdi A, Alaimo A, Etxeberria A, Fernandez-Orth J, Zamalloa T, Roura-Ferrer M, Villace P, Areso P, et al. 2011. Kv7 channels can function without constitutive calmodulin tethering. PLoS One 6:e25508. doi: 10.1371/journal.pone.0025508.

Ismailov II, Benos DJ. 1995. Effects of phosphorylation on ion channel function. Kidney Int. 48:1167–1179. doi:10.1038/ki.1995.400.

Kubisch C, Schroeder BC, Friedrich T, Lütjohann B, El-Amraoui A, Marlin S, Petit C, Jentsch TJ. 1999. KCNQ4, a novel potassium channel expressed in sensory outer hair cells, is mutated in dominant deafness. Cell 96:437–446. doi:10.1016/S0092-8674(00)80556-5.

Lerche C, Scherer CR, Seebohm G, Derst C, Wei AD, Busch AE, Steinmeyer K. 2000. Molecular cloning and functional expression of KCNQ5, a potassium channel subunit that may contribute to neuronal M-current diversity. J. Biol. Chem. 275:22395–22400. doi: 10.1074/jbc.M002378200.

Luo J-H, Fu ZY, Losi G, Kim BG, Prybylowski K, Vissel B, Vicini S. 2002. Functional expression of distinct NMDA channel subunits tagged with green fluorescent protein in hippocampal neurons in culture. Neuropharmacology 42:306–318. doi:10.1016/S0028-3908(01)00188-5.

Manville RW, Papanikolaou M, Abbott GW. 2018. Direct neurotransmitter activation of voltage-gated potassium channels. Nat. Commun. 9:1847. doi:10.1038/s41467-018-04266-w.

Martin-Batista E, Maglio LE, Armas-Capote N, Hernández G, Alvarez de la Rosa D, Giraldez T. 2021. SGK1.1 limits brain damage after status epilepticus through M current-dependent and independent mechanisms. Neurobiol. Dis. 153:105317. doi:10.1016/j.nbd.2021.105317.

Miceli F, Soldovieri MV, Joshi N, Weckhuysen S, Cooper E, Taglialatela M. 2010. KCNQ2-Related Disorders.

Michel MC, Radziszewski P, Falconer C, Marschall-Kehrel D, Blot K. 2012. Unexpected frequent hepatotoxicity of a prescription drug, flupirtine, marketed for about 30 years. Br. J. Clin. Pharmacol. 73:821–825. doi:10.1111/j.1365-2125.2011.04138.x.

Miranda P, Cadaveira-Mosquera A, González-Montelongo R, Villarroel Á, González-Hernández T, Lamas JA, Álvarez de la Rosa D, Giráldez T. 2013. The Neuronal Serum- and Glucocorticoid-Regulated Kinase 1.1 Reduces Neuronal Excitability and Protects against Seizures through Upregulation of the M-Current. J. Neurosci. 33:2684–2696. doi:10.1523/JNEUROSCI.3442-12.2013.

Nappi P, Miceli F, Soldovieri MV, Ambrosino P, Barrese V, Taglialatela M. 2020. Epileptic channelopathies caused by neuronal Kv7 (KCNQ) channel dysfunction. Pflugers Arch. Eur. J. Physiol. 472:881–898. doi:10.1007/s00424-020-02404-2.

Oliveros JC, Franch M, Tabas-Madrid D, San-León D, Montoliu L, Cubas P, Pazos F. 2016. Breaking-Cas—interactive design of guide RNAs for CRISPR-Cas experiments for ENSEMBL genomes. Nucleic Acids Res. 44:W267–W271. doi:10.1093/nar/gkw407.

Oyrer J, Maljevic S, Scheffer IE, Berkovic SF, Petrou S, Reid CA. 2018. Ion channels in genetic epilepsy: From genes and mechanisms to disease-targeted therapies. Pharmacol. Rev. 70:142–173. doi:10.1124/pr.117.014456.

Plouin P, Anderson V. 2005. Epileptic syndromes in infancy, childhood and adolescence. 4th ed. Roger, J; Bureau, M; Dravet C, editor.

Ran FA, Hsu PD, Wright J, Agarwala V, Scott DA, Zhang F. 2013. Genome engineering using the CRISPR-Cas9 system. Nat. Protoc. 8:2281–2308. doi:10.1038/nprot.2013.143.

Ronen GM, Rosales TO, Connolly M, Anderson VE, Leppert M. 1993. Seizure characteristics in chromosome 20 benign familial neonatal convulsions. Neurology 43:1355–1360.

Schroeder BC, Hechenberger M, Weinreich F, Kubisch C, Jentsch TJ. 2000. KCNQ5, a Novel Potassium Channel Broadly Expressed in Brain, Mediates M-type Currents. J. Biol. Chem. 275:24089–24095. doi: 10.1074/jbc.M003245200.

Schroeder BC, Kubisch C, Stein V, Jentsch TJ. 1998. Moderate loss of function of cyclic-AMP-modulated KCNQ2/KCNQ3 K+ channels causes epilepsy. Nature 396:687–690. doi:10.1038/25367.

Seebohm G, Strutz-Seebohm N, Baltaev R, Korniychuk G, Knirsch M, Engel J, Lang F. 2005. Regulation of KCNQ4 Potassium Channel Prepulse Dependence and Current Amplitude by SGK1 in Xenopus oocytes. Cell. Physiol. Biochem. 16:255–262. doi:10.1159/000089851.

Seebohm G, Strutz-Seebohm N, Ureche ON, Henrion U, Baltaev R, Mack AF, Korniychuk G, Steinke K, Tapken D, Pfeufer A, et al. 2008. Long QT Syndrome-Associated Mutations in KCNQ1 and KCNE1 Subunits Disrupt Normal Endosomal Recycling of I Ks Channels. Circ. Res. 103:1451–1457. doi: 10.1161/CIRCRESAHA.108.177360.

Shah MM, Mistry M, Marsh SJ, Brown DA, Delmas P. 2002. Molecular correlates of the M-current in cultured rat hippocampal neurons. J. Physiol. 544:29–37. doi: 10.1113/jphysiol.2002.028571.

Singh NA, Charlier C, Stauffer D, DuPont BR, Leach RJ, Melis R, Ronen GM, Bjerre I, Quattlebaum T, Murphy J V., et al. 1998. A novel potassium channel gene, KCNQ2, is mutated in an inherited epilepsy of newborns. Nat. Genet. 18:25–29. doi:10.1038/ng0198-25.

Singh NA, Westenskow P, Charlier C, Pappas C, Leslie J, Dillon J, Anderson VE, Sanguinetti MC, Leppert MF. 2003. KCNQ2 and KCNQ3 potassium channel genes in benign familial neonatal convulsions: Expansion of the functional and mutation spectrum. Brain 126:2726–2737. doi: 10.1093/brain/awg286.

Snyder PM, Olson DR, Kabra R, Zhou R, Steines JC. 2004. cAMP and serum and glucocorticoid-inducible kinase (SGK) regulate the epithelial Na+ channel through convergent phosphorylation of Nedd4-2. J. Biol. Chem. 279:45753–45758. doi:10.1074/jbc.M407858200.

Soldovieri MV, Boutry-Kryza N, Milh M, Doummar D, Heron B, Bourel E, Ambrosino P, Miceli F, De Maria M, Dorison N, et al. 2014. Novel KCNQ2 and KCNQ3 mutations in a large cohort of families with benign neonatal epilepsy: First evidence for an altered channel regulation by syntaxin-1A. Hum. Mutat. 35:356–367. doi:10.1002/humu.22500.

Surti TS, Huang L, Jan YN, Jan LY, Cooper EC. 2005. Identification by mass spectrometry and functional characterization of two phosphorylation sites of KCNQ2/KCNQ3 channels. Proc. Natl. Acad. Sci. U. S. A. 102:17828–17833. doi:10.1073/pnas.0509122102.

Surur AS, Bock C, Beirow K, Wurm K, Schulig L, Kindermann MK, Siegmund W, Bednarski PJ, Link A. 2019. Flupirtine and retigabine as templates for ligand-based drug design of K V 7.2/3 activators. Org. Biomol. Chem. 17:4512–4522. doi:10.1039/c9ob00511k.

Wang HS, McKinnon D. 1995. Potassium currents in rat prevertebral and paravertebral sympathetic neurones: control of firing properties. J. Physiol. 485:319–335. doi: 10.1113/jphysiol.1995.sp020732.

Wang HS, Pan Z, Shi W, Brown BS, Wymore RS, Cohen IS, Dixon JE, McKinnon D. 1998. KCNQ2 and KCNQ3 potassium channel subunits: Molecular correlates of the M-channel. Science (80-.). 282:1890–1893. doi:10.1126/science.282.5395.1890.

Wesch D, Miranda P, Afonso-Oramas D, Althaus M, Castro-Hernández J, Dominguez J, Morty RE, Clauss W, González-Hernández T, Alvarez De la Rosa D, et al. 2010. The neuronal-specific SGK1.1 kinase regulates δ-epithelial Na+ channel independently of PY motifs and couples it to phospholipase C signaling. Am. J. Cell Physiol. 299:779–790. doi: 10.1152/ajpcell.00184.2010.

Wickenden AD, Roeloffs R, McNaughton-Smith G, Rigdon GC. 2004. KCNQ potassium channels: Drug targets for the treatment pf epilepsy and pain. Expert Opin. Ther. Pat. 14:457–469. doi:10.1517/13543776.14.4.457.

Wickenden AD, Zou A, Wagoner PK, Jegla T. 2001. Characterization of KCNQ5/Q3 potassium channels expressed in mammalian cells. Br. J. Pharmacol. 132:381–384. doi: 10.1038/sj.bjp.0703861.

Wulff H, Castle NA, Pardo LA. 2009. Voltage-gated potassium channels as therapeutic targets. Nat. Rev. Drug Discov. 8:982–1001. doi:10.1038/nrd2983.

Xiong Q, Sun H, Zhang Y, Nan F, Li M. 2008. Combinatorial augmentation of voltage-gated KCNQ potassium channels by chemical openers. Proc. Natl. Acad. Sci. 105:3128–3133. doi:10.1073/pnas.0712256105.

