## Supplementary Info for "Activation of SGK1.1 up-regulates the M-current in the presence of epilepsy mutations"

**This PDF file includes:**

Figures S1 to S3

SUPPLEMENTARY FIGURE 1

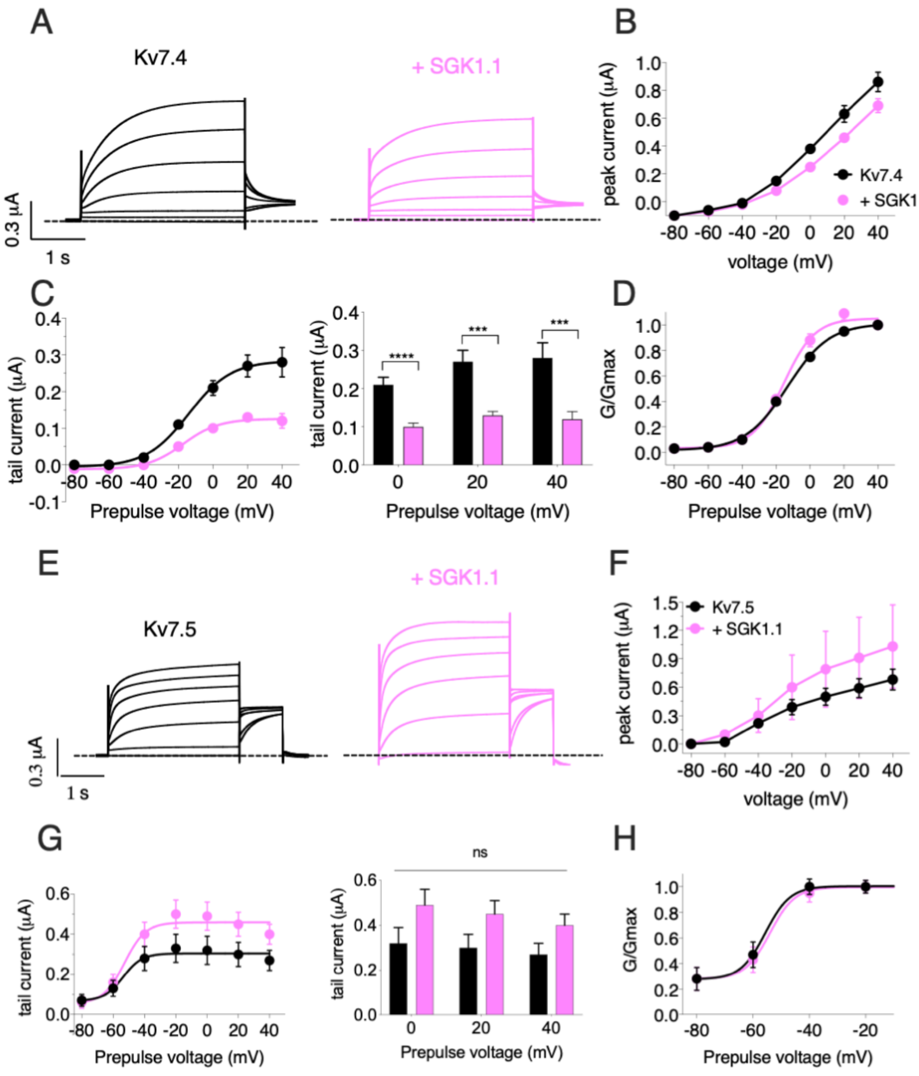


**Figure S1. SGK1.1 fails to increase homomeric Kv7.4 and Kv7.5 currents, similarly to previously reported results for homomeric Kv7.2 and Kv7.3**. (A) Currents elicited in *Xenopus* oocytes after coinjection cDNAs from Kv7.4 channel alone or in combination with wild-type SGK1.1. (B) Peak current/voltage relationship. (C) Tail current (left) and tail currents measured at -30 mV after 0, +20 or +40 mV depolarizing pulses for the indicated construct combinations (right). Values represent mean ± SEM (Multiple t-test Holm-Sidak correction method;***p<0.0005, ****p<0.0001). (D) Normalized conductance. (E) Currents elicited in *Xenopus* oocytes after coinjection of Kv7.5 channel alone or in combination with wild-type SGK1.1. (F) Peak current/voltage relationship. (G) Tail current (left) and tail currents measured at -30 mV after 0, +20 or +40 mV depolarizing pulses for the indicated construct combinations (right). Values represent mean ± SEM (Multiple t-test Holm-Sidak correction method; ns, not significant). (H) Normalized conductance.

SUPPLEMENTARY FIGURE 2

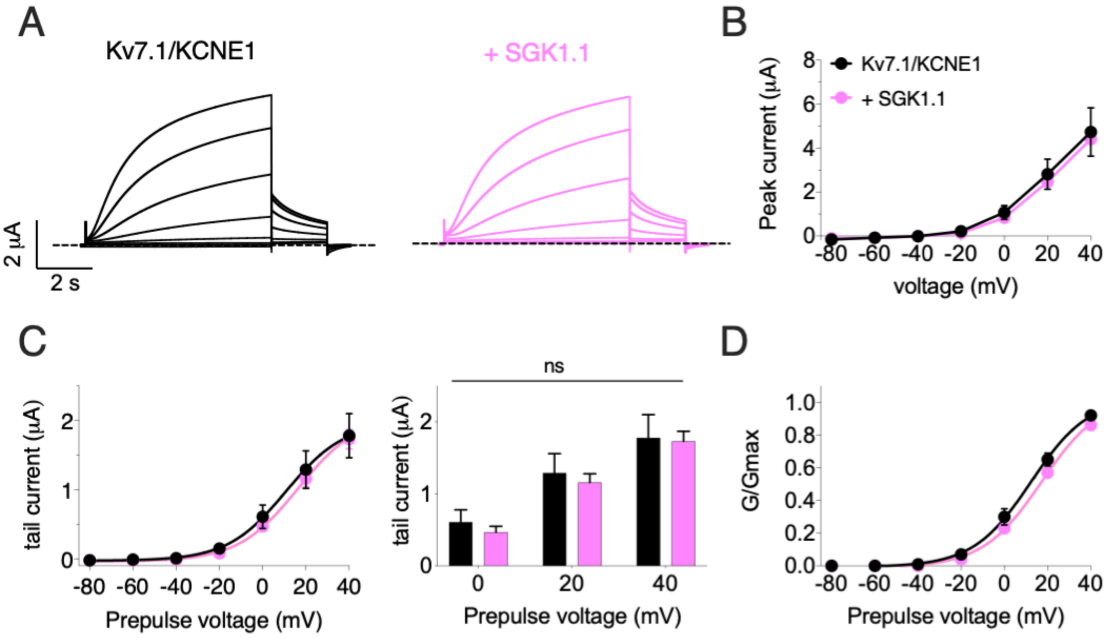


**Figure S2. SGK1.1 does not affect the major repolarizing cardiac current Kv7.1/KCNE1 ($I_{KS}$).** (A) Current elicited in Xenopus oocytes after coinjection of cDNAs from Kv7.1/KCNE1 channel alone (first panel) or in combination with wild-type SGK1.1 (second panel). (B) Peak current/voltage relationship. (C) Tail current (left) and tail currents measured at -30 mV after 0, +20 or +40 mV depolarizing pulses for the indicated construct combinations (right). Values represent mean ± SEM (Multiple t-test Holm-Sidak correction method; ns, not significant). (D) Normalized conductance.

SUPPLEMENTARY FIGURE 3

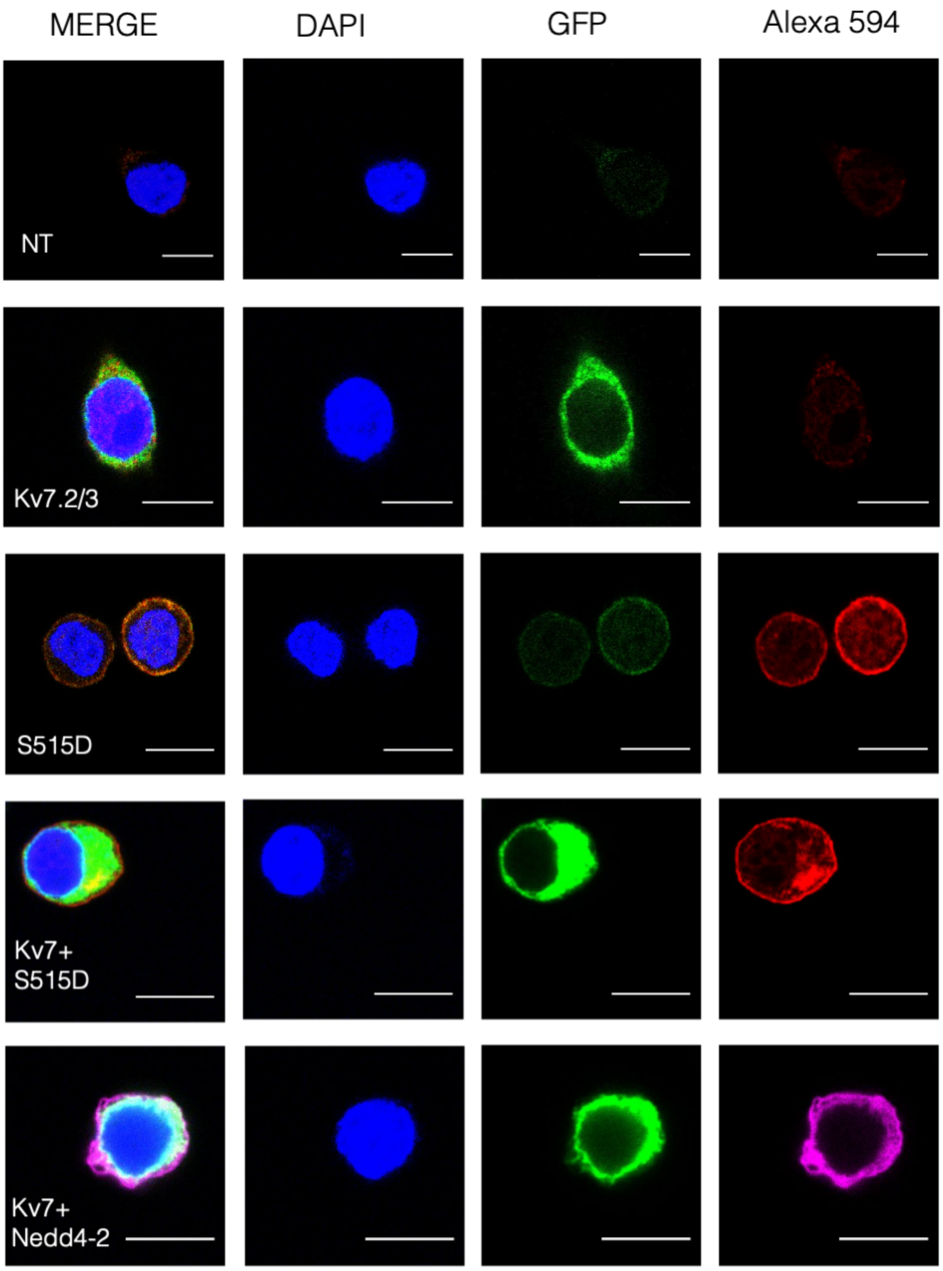


Figure S3. Representative confocal images showing expression of Kv7.2/3, SGK1.1 and Nedd4-2. *Sgk1*-KO N2a cells were transfected with the indicated constructs or left untransfected (NT). Twenty-four hours after transfection, cells were permeabilized and incubated with specific antibodies. Cells were stained with DAPI for nuclei detection. Transfection with Kv7 subunits was detected as green fluorescence as constructs were fused to YFP. SGK1.1 was detected using a rabbit polyclonal antibody and a secondary anti-rabbit antibody conjugated to Alexa Fluor® 594. Nedd4-2 expression was detected with anti-Nedd4-2 and Alexa Fluor® 594 fused anti-rabbit. To facilitate identification, Nedd4-2 signal is shown in magenta. Bars correspond to 10 μM.
